# Profiling Glioma Stem Cell Dynamics via 3D-based Cell Cycle Reporter Assays

**DOI:** 10.1101/2024.02.16.580653

**Authors:** Dorota Lubanska, Antonio Roye-Azar, Sami Alrashed, Alan Cieslukowski, Mohamed Soliman, Ana C. deCarvalho, Abdalla Shamisa, Swati Kulkarni, Lisa A. Porter

## Abstract

Successful containment of unwanted cell cycle progression in tumours such as glioblastoma (GBM) requires targeted therapeutic approaches which rely on understanding cell cycle dynamics in response to microenvironmental stimuli. Glioma Stem Cells (GSCs) can drive tumour initiation, recurrence, therapy resistance, and are often attributed to the heterogeneity and plasticity of GBM. *In vitro* models using patient-derived GSCs provide a life relevant tool for exploration of complex molecular mechanisms underlying the aggressive characteristics of GBM. Introduction of 3D tissue culture systems permits the study of spatial complexity of the tumour mass and enables control over diverse conditions within the surrounding microenvironment. This chapter demonstrates detailed methods to study spatio-temporal changes to the cell cycle dynamics using available fluorescent cell cycle reporter systems in combination with bioinformatics-based signal intensity and localization analysis. We present a successful approach that investigates the 3D cell cycle dynamics of GSC populations. This approach utilizes GBM neurosphere and organoid cultures, which are assessed over time and under therapeutic pressure. These models can be further explored, manipulated, and customized to serve specific experimental designs.

## Introduction

Despite significant advancements in our understanding of brain cancer biology, cancer stem cells remain a critical obstacle to the eradication of tumours such as glioblastoma (GBM) ^1,2^. Glioma stem cells (GSCs) share multiple characteristics with normal neural stem cells including self-renewal and multipotent differentiation and are a potential source of GBM initiation and recurrence ^3,4^. Diverse characteristics of GSCs have been explored over the years, focusing on their response to DNA damage, evasion of apoptosis and lineage plasticity which contribute to radio- and chemo-resistance, including the standard of care temozolomide ^3,5,6,7,8^.

Mechanistically, regulation over the cell cycle machinery plays a pivotal role in complex fate decisions dictating proliferation/expansion, quiescence (arrested growth) and differentiation of all forms of stem cells, including GSCs. The proliferation kinetics of cancer stem cells often differ from those of other cells within the tumour and can range from being fast-cycling, slow-cyclin or quiescent ^9,10^. In GBM there is a need to characterize how these properties relate to invasiveness of the tumour ^11^ and to consider the consequence of targeting cycling GSCs on the long-term therapy response of the tumour ^12^.

Critical molecular players possess the capacity to control the cell cycle and maintain GCSs in G1 phase, leading to slow proliferation or cell cycle exit ^13^. Multiple studies have demonstrated that cancer stem cells can rapidly proliferate upon the application of growth factors and mechanical changes to the microenvironment *in vitro* ^14,15^. Conversely, spatial changes can also prompt quiescence ^16^, further highlighting the essential role of the surrounding microenvironment in regulating the proliferation kinetics of cancer stem cells. The evolution of 3D *in vitro* systems with well-defined and modifiable microenvironment has added to the repertoire of tools available to study the spatial and temporal changes of select cells within a tumour ^17,18,19^. 3D culturing of tumour cells or tissue fragments *in vitro* can be broadly divided into two types of cultures: neurospheres and tumour organoids, sometimes referred to as spheroids, each with their own advantages and disadvantages.

Cells grown in well characterized suspension cultures with nutrient-rich medium, known as neurosphere cultures, offer the advantage of a 3D culture context, facilitate the selection of GSCs, are easily reproducible and relatively efficient. They do however have a significant limitation, they lack microenvironment, and consequently, are not fully representative of systems that allow cells to interact with a niche or receive signaling from surrounding extracellular matrix or cellular components of the stroma.

Tumour organoids, or spheroids, are self-organizing 3D microtissues that effectively recapitulate characteristics of tumour growth and allow for easy manipulation of the tumour cells. They also enable alteration of the composition and physical properties of the extracellular matrix with high fidelity. Many diverse studies have used tumour organoid models to examine the influence of the microenvironment on GSC behaviour, including invasiveness, heterogeneity, and proliferation ^20,21^. Diverse properties of the microenvironment ranging from physical rigidity, oxygen and nutrient levels and response to other cell types including those of the immune system have been studied. These further permit manipulation of molecular mechanisms and direct observation of effects on GBM progression and aggressiveness, aspects which are limited in *in vivo* models.

In the cell cycle field, there are a plethora of powerful cell cycle reporters available which allow for the study of spatial and temporal factors impacting the cell cycle during tumour development. Reporters which provide cell cycle phase information facilitate the study of the dynamics within a heterogeneous tumour mass faced with different conditions or stressors, including those occurring during treatment pressure. **F**luorescence **u**biquitin **c**ell **c**ycle **i**ndicator (FUCCI) is based on the mutually exclusive activity of two proteins, cdt1 and geminin, in the G0/G1 and S/G2/M phases of the cell cycle, respectively ^22^. The proteins in the FUCCI system are developed as fluorescently labeled probes which can determine the cell cycle phase or transition in individual cells based on the quantified fluorescent signal of the nucleus. This allows for the exploration of cell cycle dynamics in live cell and *in vivo* assays. **D**NA **H**elicase **B** (DHB)-based CDK2 activity reporter uses established CDK2 phosphorylation sites on a fragment of fluorescently labeled human Helicase B ^23^. The subcellular localization of DHB serves as a CDK2 activity sensor; DHB localizes to the nucleus in its unphosphorylated state while it is exported to the cytoplasm upon phosphorylation. The fluorescent signal is localized and scored via a bioinformatics-based double masking quantification approach. The Cytoplasm/Nucleus signal ratio is then used to determine the levels of CDK2 activity in an individual cell.

The protocols described in this chapter include the generation, culture, and manipulation of patient derived GSC neurospheres and organoids as well as methods of imaging and analysis which have been optimized in our lab using several individual patient-derived GSC populations. We demonstrate detailed techniques and the steps required to generate successful GSC-based 3D *in vitro* models that report cell cycle changes in a spatial and temporal manner. Additionally, these methods can be further modified and customized to align with the requirements of specific experimental designs, thereby addressing particular research questions.

## 2. Materials

### 2.1. Patient-derived GSC culture

1. Primary glioblastoma cells (identified as: HF3253 and WRH005) were obtained through enzymatic extraction of GBM tissues derived at surgery.
2. GSC media:

a. Dulbecco’s Modification of Eagle’s Medium (DMEM)/ Ham’s F12 50/50 Mix.
b. N-2 Supplement (100X).
c. Epidermal Growth Factor (EGF) Recombinant Human Protein.
d. Fibroblast Growth Factor (FGF) – Basic Human, Recombinant expressed in *E. Coli*.
e. Gentamicin solution.
f. Antibiotic-Antimycotic (100X).
3. Cell culture grade Bovine Serum Albumin (BSA), heat shock fraction, protease free, low endotoxin.
4. Ca^2+^/ Mg^2+^ - free PBS
5. T25 regular tissue culture flasks with red vented caps (Sarstedt).

### 2.2. Lentiviral transduction

1. FUCCI or DHB-mVenus Lentiviral particles generated using pBOB-EF1-FUCCI-Puro plasmid (#86849, Addgene) or DHB-mVenus (#136461, Addgene).
2. Dulbecco’s Modification of Eagle’s Medium (DMEM)/ Ham’s F12 50/50 Mix.
3. Polybrene 10mg/ml stock.
4. Puromycin 1mg/ml stock.
5. 96-well polystyrene tissue culture dishes.

### 2.3. Fluorescent Activated Cell Sorting (FACS) enrichment

1. PBS with 2mM EDTA
2. Flowcytometry tubes

### 2.4. Organoid generation and culture

1. Media composition same as for GSC culture.
2. Matrigel
3. Parafilm
4. 96-well U-bottom ultra-low attachment (ULA) plates (Greiner)
5. 24-well ULA plates

### 2.5. Sectioning

1. 30% sucrose solution in PBS.
2. 30% sucrose solution in 4% PFA (PBS based).
3. Optimal Cutting Temperature (OCT) cryoembedding matrix
4. 1.5 mL Eppendorf tubes.
5. Charged tissue slides and coverslips.
6. Fluorescence microscopy mounting medium.
7. 1μg/mL of Hoechst 33342 counterstain in Hank’s Balanced Salt Solution.

### 2.6. SA-**β**-Gal staining

1. X-gal Stock: 40 mg/ml in dimethylformamide, store at –20°C protected from light.
2. SA-β-gal Staining Solution (minus X-gal; 100 mL): 0.221g of Potassium Ferrocyanide (MW 422.4 g/mol; 5 mM final), 0.165 g of Potassium Ferricyanide (MW 329.2 g/mol; 5 mM final), 200 mL of 1 M MgCl2 (2 mM final), pH6.0
3. Complete staining solution: Add 25 ml of 40 mg/ml X-gal per mL of staining solution (1 mg/mL final)

## 3. Methods

### 3.1. Preparation of GBM patient-derived GSC cultures

1. Cells are cultured as free-floating spheres in DMEM/F-12 50/50 culture media supplemented with 1X N-2 supplement, 20ng/mL EGF, 10ng/mL FGF, 50 µg/mL gentamicin solution, 1X antibiotic-antimycotic, and 0.5mg/mL BSA in T25 flasks at 37°C and 5% CO_2_ seeded at 10x10^5^ cells per 1mL of media for 5-6 days or till they reach size of 250mm in diameter.
2. Cells successfully subcultured as neurospheres to at least passage 4 are used in assays. (*see* **Note 1**)
3. Contents of the flask are transferred to a 15mL conical tube and allow to sediment by gravity.
4. The supernatant is removed, and spheres are suspended and incubated in 2mL of Ca^2+^/ Mg^2+^ - free PBS solution for 15min at room temperature with gentle pipetting every 5 minutes till homogenous single cell suspension is achieved.
5. The cell suspension is centrifuged at 800x g for 5 minutes at 4°C.
6. The cells are resuspended in 2mL of growth media.
7. The cells are counted via trypan blue exclusion assay using hemocytometer or automatic cell counter and live cell number is used in the following applications.

### 3.2. Lentiviral transduction and selection of transduced GSCs

1. Primary GSCs are suspended in DMEM/F-12 50/50 culture media containing 8μg/mL polybrene at 10^5^ cells/mL. (*see* **Note 2**)
2. The suspension is transferred to 96-well U-bottom shaped ULA plate at 100μL/well.
3. Lentivirus (FUCCI or DHB-mVenus) is added to the wells at MOI of 2. (*see* **Note 3**)
4. Lentiviral media is removed and replaced with 125 μL of growth media per well. (*see* **Note 4**)
5. FUCCI transduced GSCs are selected by treatment with 1μg/ml of Puromycin at 48 hours post transduction.
6. DHB-mVenus transduced GSCs are suspended in PBS/2mM EDTA at 10^6^/mL in flowcytometry tubes and sorted using FACS.
7. Cells are incubated at 37°C and 5% CO_2_ to form spheres for 5 days with addition of 50 μL of fresh media on day 3. (*see* **Note 5**)

### 3.3. Generation of organoids

1. Transduced spheres are pipetted in 10 μL of media onto the 10cmx10cm square of parafilm placed inside 150mm dish, 1 cm apart.
2. The excess of media is gently pipetted off the spheres.
3. Matrigel, defrosted overnight, is added as a 20 μL droplet to each sphere.
4. The dish containing parafilm with embedded spheres is placed inside an incubator at 37°C and 5% CO_2_ for 10 minutes.
5. Embedded spheres are then transferred using a P200 pipette tip to 24-well ULA plate containing 1mL of prewarmed growth media, one sphere per well.
6. Organoids are allowed to form in the stationary phase for 7 days.
7. The plate is then moved onto a shaker set to low speed and organoid formation proceeds with nutrient circulation.

### 3.4. Organoid sectioning

1. Collected organoids are fixed in 30% sucrose in PFA overnight at 4°C.
2. Fixed samples are then moved to 30% sucrose and stored at 4°C.
3. Prior to cryosectioning the organoids are transferred to Eppendorf tubes and submerged in 0.5 mL of OCT matrix.
4. The tubes are placed inside a vacuum oven and incubated at 15-20 psi for 15 minutes.
5. The organoids are prepared for cryosectioning on OCT covered chucks and incubated inside the cryostat for 20 minutes.
6. The sections are cut at 10 μm setting and transferred onto the prepared glass slides. (*see* **Note 6**)
7. Organoid sections on slides are stored at -20°C.

### 3.5. SA-**β**-Gal staining

1. Slides with organoid sections are defrosted for 2 minutes at room temperature.
2. The slides are then washed 3x 2 minutes with PBS and 1μg/mL of Hoechst counterstain is applied for 15 minutes for DHB-mVenus organoid sections)
3. The sections are then submerged in complete SA-b-Gal solution by placing the slides in plastic slide holder.
4. The sections are incubated overnight in an oven at 37°C, protected from light.
5. Slides are washed with distilled water and mounting media is applied to the sections followed by a coverslip application.

### 3.6. Imaging & analysis

1. Spheres and organoids collected for imaging are cultured in phenol free based GSC media. (*see* **Note 7 & 8**)
2. The samples are imaged using BioTek Cytation5 imager and GEN5 software in Z-stack imaging mode.
3. The images are processed via Z-projection and Focus stacking method.
4. Images are exported in jpeg format.
5. The images are analyzed using ImageJ software.
6. In tab Image select 8-bit option the image type.
7. Using Adjust selection in the Image tab go to Threshold.
8. Adjust the threshold to match the signal in the image and select Apply.
9. In the Analyze tab, select Set Measurements.
10. In the Measurements selection, select Integrated Density Values (IDV)and Limit to threshold. Apply changes.
11. Select Measure in the Analyze tab to obtain IDV which corresponds to fluorescence intensity of the analyzed image.

### 3.7. Double masking analysis

1. CellProfiler 4.2.5 software is used for the analysis (www.cellprofiler.org). The left side of the GUI is split into two sections: the pre-set input modules “Images”, “Metadata”, “Names And Types”, and “Groups”; and the empty pipeline panel where modules can be added.
2. To add modules, right click the pipeline panel to select which modules to add to the pipeline. The step-by-step panel settings for each module are available in *Supplementary materials*.
3. The modules are used in the following order:

a. Images: used to pass all specified files to downstream modules. The files are fluorescent microscopy TIFF images, without size annotations or labels, of DHB-mVenus expressing cells (GFP channel) counterstained with DAPI for nuclei.
b. Names And Types: assigns a meaningful name to each image by which other modules will refer to it.
c. Correct Illumination Calculate: calculates an illumination function that is used to correct uneven illumination/lighting/shading or to reduce uneven background in images. This calculates the illumination function for the DAPI channel images.
d. Correct Illumination Apply: applies the illumination function created in the previous step to correct the DAPI channel images.
e. Correct Illumination Calculate: calculates the illumination function for the GFP channel images.
f. Correct Illumination Apply: applies the illumination function created in the previous step to correct the GFP channel images.
g. Identify Primary Objects: identifies biological objects of interest and requires grayscale images containing bright objects on a dark background. The nuclei are identified as the primary objects.
h. Edit Objects Manually: creates, removes, and edits objects previously defined. If the noise around the corners of the images worsens the segmentation in those areas, the nuclei in those areas are deleted.
i. Identify Secondary Objects: identifies objects using other objects as a starting point. The cytoplasms of individual cells are identified as secondary objects using their corresponding nuclei.
j. Measure Object Size Shape: measures several area and shape features of identified objects. Several area and shape features are taken of each individual cytoplasm and nuclei.
k. Calculate Math: takes measurements produced by previous modules and performs basic arithmetic operations. The area of the cytoplasm objects is divided by the area of their corresponding nucleus objects.
l. Filter Objects: removes selected objects based on measurements produced by another module. The nuclei are filtered according to the area covered by their corresponding individual cytoplasm. The nuclear area had to be at least 1.5x smaller than the cytoplasmic area or else it was discarded.
m. Filter Objects: individual cytoplasms are filtered according to the area they covered in respect to their nuclei. The cytoplasmic area had to be at least 1.5x larger than the nuclear area or else it was discarded.
n. Relate Objects: assigns a relationship to objects so that nuclei can be grouped with their cytoplasm.
o. Mask Objects: removes objects outside of a specified region. The nuclei mask is used in conjunction with the cytoplasm mask to create a mask that contains the cytoplasm excluding the area covered by the nuclei.
p. Measure Object Intensity: measures several intensity features for the identified objects. These measurements are taken using the filtered cytoplasmic (excluding nuclei) and filtered nuclei masks on the GFP channel images.
q. Measure Object Intensity Distribution: measures the spatial distribution of intensities within each object. These measurements are taken using the filtered cytoplasmic (excluding nuclei) and filtered nuclei masks on the GFP channel images.
r. Calculate Math: The Integrated Intensity of the cytoplasm is divided by the integrated intensity of the nucleus.
s. Measure Object Size Shape: Several area and shape features are taken of the filtered cytoplasmic (excluding nuclei) and filtered nuclei masks.
t. Export To Spreadsheet: exports measurements into one or more files that can be opened in Microsoft Excel or other spreadsheet programs.

## Figure captions

**Figure 1.**
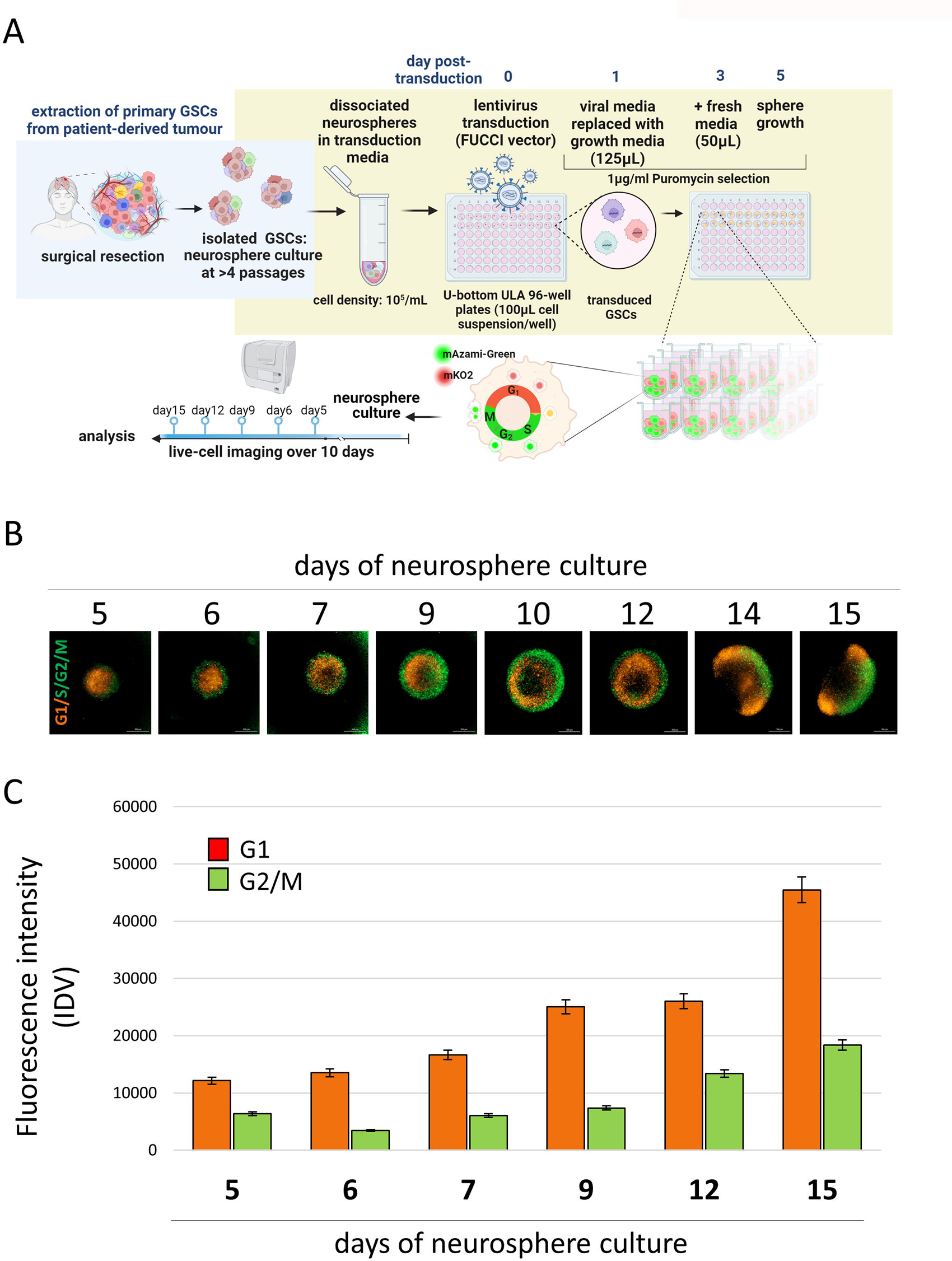
Spatio-temporal analysis of cell cycle phase changes in FUCCI-transduced neurospheres derived from patient GSCs. A. Schematic of the assay workflow. Created with BioRender.com. B. 3D Z-stack projection-derived images of GBM neurospheres cultured over a 15-day timecourse. Images taken at 24-hour intervals. C. Quantified fluorescence signal of the respective cell cycle phases. Integrated density values represent cells assigned to G1 (orange-red) versus S/G2/M (green) phase by FUCCI reporter. The signal is corrected by the diameter of analyzed neurospheres. The cells demonstrated distinct spatial cell cycle dynamics over time. Cell culture conditions caused accumulation of cells in G1 phase of the cell cycle.

**Figure 2.**
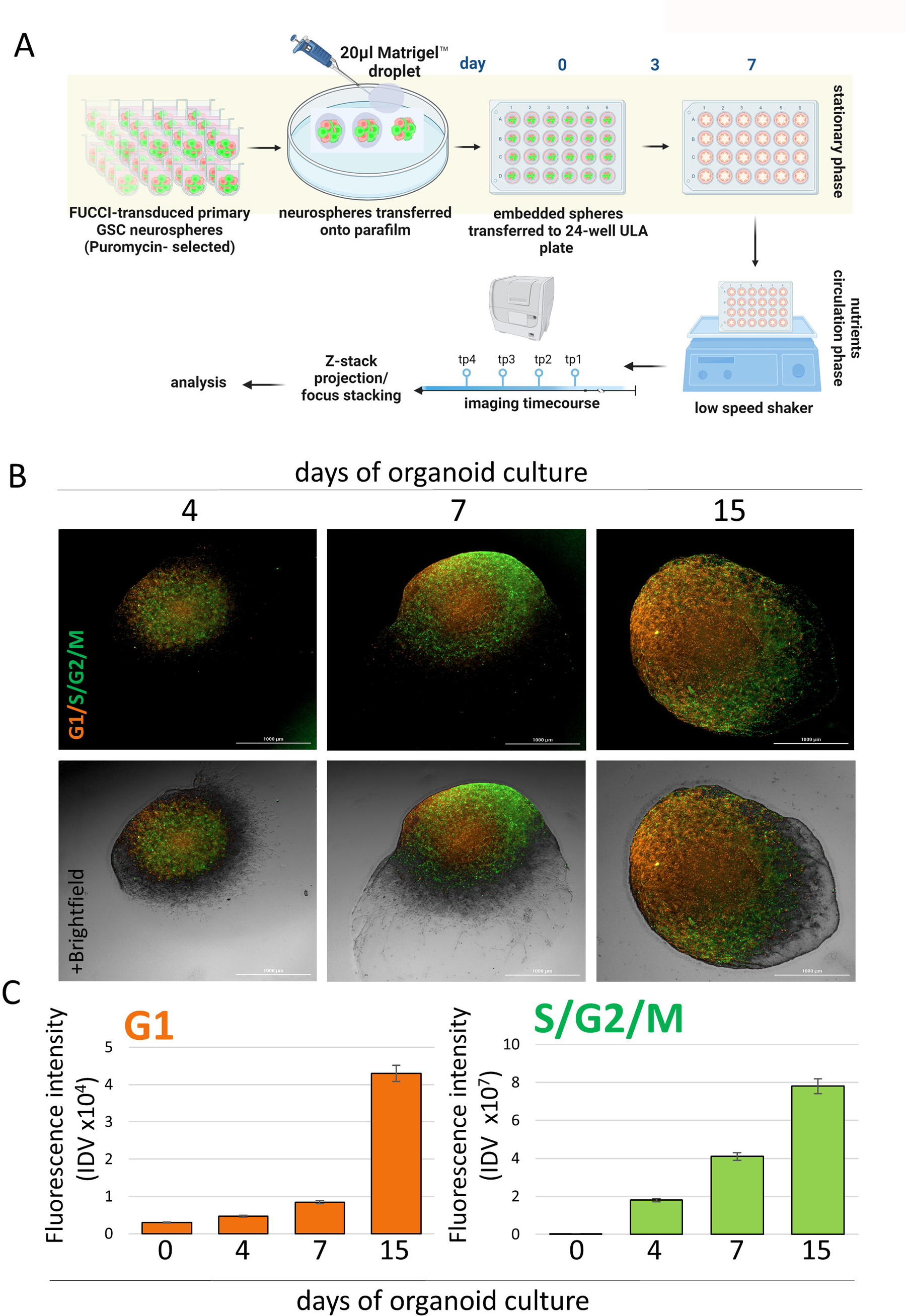
Analysis of the microenvironment-mediated changes of the spatio-temporal dynamics of GSC populations in the setting of the FUCCI brain tumour organoid model. A. Schematic of the assay workflow. Created with BioRender.com. B. GBM patient derived FUCCI organoids imaged over time at day 4, 7 and 15 using vis Z-stack projection. Fluorescent images only-top panel, overlay of fluorescent images over brightfield-bottom panel. C. Quantified fluorescence signal of the respective cell cycle phases reported by FUCCI. Values graphed as integrated density for cells assigned to G1 phase (orange-red) versus S/G2/M cell cycle phase (green). In the setting of the applied microenvironment GSCs display distinct spatial patterns of cell cycle phases over time and cells accumulate primarily in green S/G2/M (S/G2/M IDV of 7.8x10^7^ vs. G1 IDV of 4.3x10^4^ at day 15).

**Figure 3.**
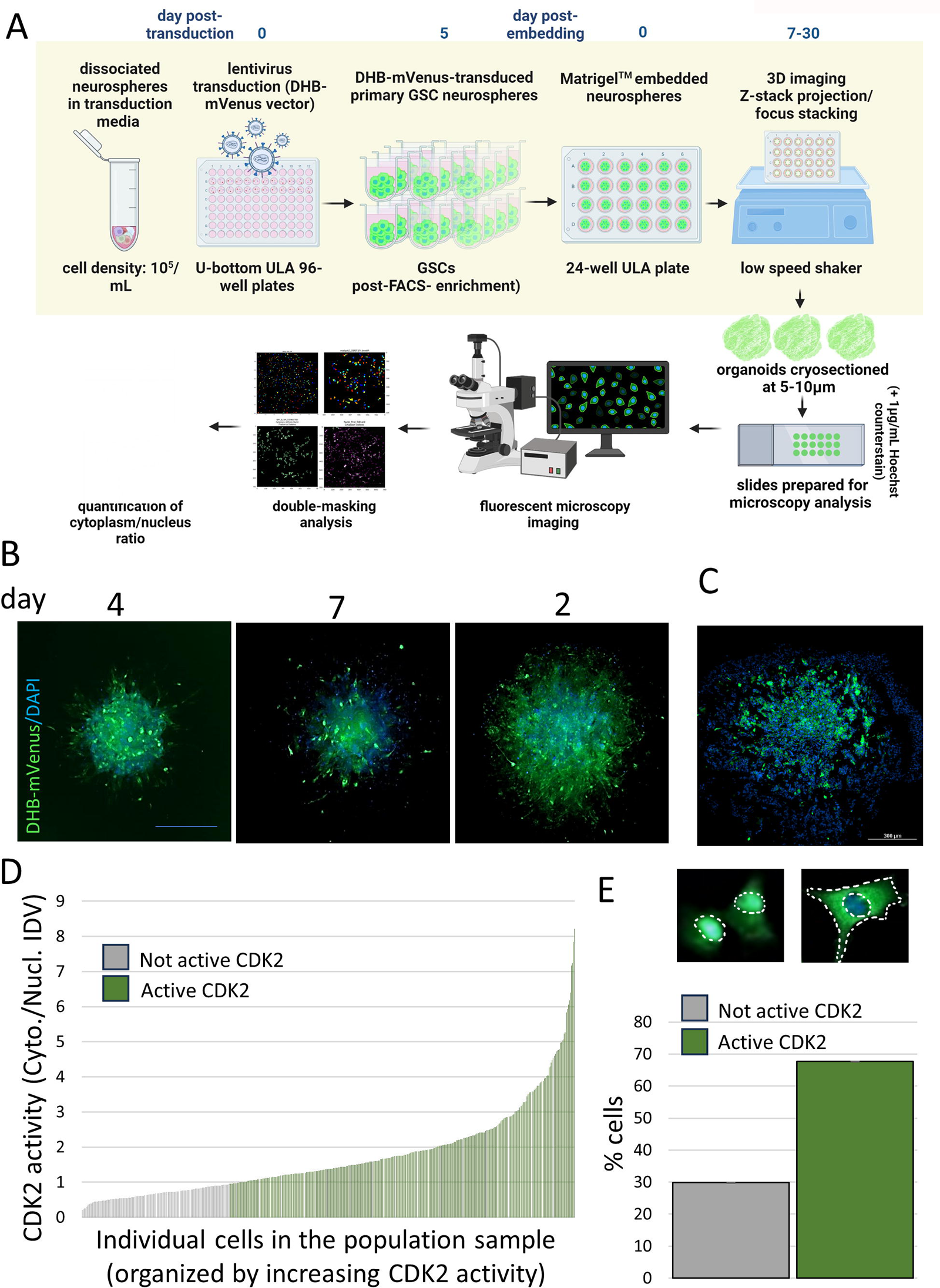
Generation and analysis of GBM patient DHB-mVenus CDK2 reporter organoids. A. Schematic of the workflow. Created with BioRender.com. B. Organoids imaged using Z-stacking mode at 4, 7, 22 days of culture. C. A representative image of a cryosection of DHB-mVenus CDK2 reporter organoid collected on day 22 of culture. D. DHB-mVenus+ scored for CDK2 activity levels using reporter-based fluorescence and double masking method. The graph demonstrates the range of CDK2 activity levels over the population analyzed. E. Number of cells expressing active CDK2, not active CDK2 or CDK2 of intermediate activity quantified and graphed as percentage of the analyzed population. DHB mVenus-transduced GSCs show successful expression of the reporter over duration of the organoid culture and the signal is preserved in fixed/processed and sectioned organoid microtissues. Double masking modality approach allows to adequately assess the levels of fluorescent signal of subcellular localization.

**Figure 4.**
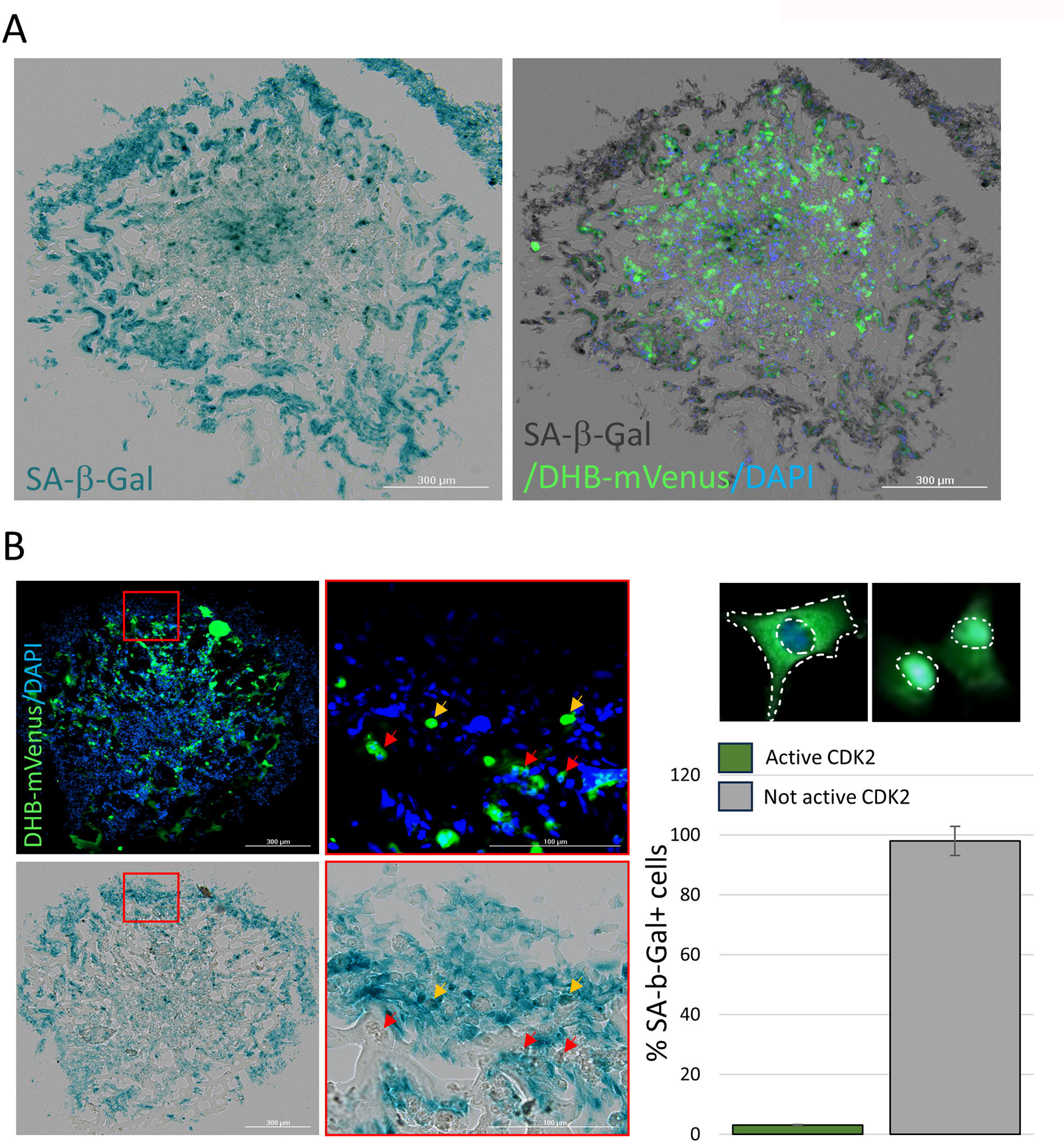
DHB-mVenus organoids stained for SA-β-Gal activity. A. X-gal stain (left) overlaid onto DHB-mVenus fluorescence in organoid-derived cryosections at day 22. B. High-magnification images of the DHB-mVenus cryosections stained for SA-β-Gal activity. C. Quantified % of SA-β-Gal activity+ cells in CDK2 active cells (red arrows) and CDK2 inactive cells (yellow arrows). Combined analysis of the reporter signal with a senescence marker demonstrates exclusive activity of CDK2 and SA-β-Gal.

## Notes

1. Selection of GSCs from tumour tissue extractions takes place on regular tissue culture surface. Only cells forming spheres are collected leaving behind any stromal cell component which remains attached to the bottom of the dish.
2. GSC cultures for designated viral transduction should be above 90% of viability for best results.
3. Primary GSC cultures drastically vary in sensitivity to lentiviral transduction. It is recommended that individual MOI is tested and validated for each culture to minimize cell death.
4. If nuclear counterstain is required at the time of 3D imaging 1μg/mL Hoechst can be added to the media at 24h post transduction and later to media hosting organoids.
5. Neurospheres above 150μm in diameter are best suited for observation of spatial changes in fluorescence marking population dynamics. The timing of each step specified in the protocol produces desired results for majority of the cultures, however, it might require adjustment for some cell lines.
6. Organoid sections between 5-10 μm are recommended for proper analysis of DHB-mVenus signal localization.
7. Aspirating media off the well during the early stages of organoid culture improves efficiency and quality of organoid imaging. Attention must be paid to prevent organoid drying.
8. The 3D imaging sessions of the organoids through Z-stacking may take significant amount of time, it is recommended the organoids are imaged at 37°C and 5% CO_2._
9. Double masking analysis pipeline files are available in *Supplementary materials*.
10. To facilitate cell segmentation and double masking, the confluency of the fluorescently labelled cells should be low enough to ensure some space between cells. Additionally, the cells should be evenly stained. There are 4 cases which might result in outliers in the cytoplasmic/nucleus ratio:
10.1. The integrated intensity of the nucleus is very low. This would mean that the nucleus was not stained well enough to be properly segmented or that a nucleus was detected next to a cell and was accidentally grouped with a cytoplasm that it shouldn’t have been grouped with.
10.2. The integrated intensity of the nucleus is too high. Attention must be paid to make sure that none of the nuclei were clumped so if a reading is very high then this represents a normal variation in the population where the nucleus is large i.e. not a true outlier.
10.3. The integrated intensity of the cytoplasm is very low. This would mean that the cytoplasm was too small/not stained well enough to be properly segmented.
10.4. The integrated intensity of the cytoplasm is too high. The program is accurate at segmenting the cytoplasm of the individual cells so if a reading is very high then this represents a normal variation in the population where the cytoplasm is large i.e. not a true outlier.

