## Supplemental material for "Profiling Glioma Stem Cell Dynamics via 3D-based Cell Cycle Reporter Assays"

**Supplementary materials**

Module panel settings

CellProfiler 4.2.5 was used for this analysis, please refer to www.cellprofiler.org.

The left side of the GUI is split into two sections: the pre-set input modules “Images”, “Metadata”, “NamesAndTypes”, and “Groups”; and the empty pipeline panel where modules can be added. To add modules right click the pipeline panel to select which modules to add to the pipeline. The modules used, their order, and their rationale is as follows:

**Images**

This module is used to pass all specified files in the file list to downstream modules.

The images were dropped into the files panel. The images used were fluorescent microscopy images (without size annotations or labels) of nuclei stained with DAPI, cytoplasms stained with GFP, and DHB-Ven green CDK2 sensor.

**Names And Types**

This module assigns a meaningful name to each image by which other modules will refer to it. “Add another image” was clicked to add a second criteria panel to organize the images. The panel settings are as follows:

- Assign a name to **Images matching rules**
- Match **All** of the following rules
- **File Does Contain DAPI**
- Name to assign these images **DAPI**
- Select the image type **Greyscale image**
- Set intensity range from **Image metadata**
- Assign a name to **Images matching rules**
- Match **All** of the following rules
- **File Does Contain GFP**
- Name to assign these images **GFP**
- Select the image type **Greyscale image**
- Set intensity range from **Image metadata**

**CorrectIlluminationCalculate**

This module calculates an illumination function that is used to correct uneven illumination/lighting/shading or to reduce uneven background in images. This calculates the illumination function for the DAPI channel images. The panel settings are as follows:

- Select the input image: **DAPI**
- Name the output image: **DAPI_ILLUM**
- Select how the illumination function is calculated: **Background**
- Block size: **20**
- Rescale the illumination function?: **No**
- Calculate function for each image individually, or based on all images?: **Each**
- Smoothing method: **Median Filter**
- Method to calculate smoothing filter size: **Automatic**

The block size chosen is important and should be adjusted to the objects in the image. The block size should be large enough to cover several objects and include some background but small enough that it won’t overlook significant local variations in illumination. Artifacts may be introduced into the image with an inappropriate block size.

**CorrectIlluminationApply**

This module applies the illumination function created in the previous step to correct the DAPI channel images. The panel settings are as follows:

- Select the input image **DAPI_ILLUM**
- Name the output image **DAPI_ILLUM_CORRECTED**
- Select the illumination function **DAPI_ILLUM**
- Select how the illumination function is applied **Subtract**

**CorrectIlluminationCalculate**

This module calculates the illumination function for the GFP channel images. The panel settings are as follows:

- Select the input image **GFP**
- Name the output image **GFP_ILLUM**
- Select how the illumination function is calculated **Background**
- Block size **22**
- Rescale the illumination function? **No**
- Calculate function for each image individually, or based on all images? **Each**
- Smoothing method **Median Filter**
- Method to calculate smoothing filter size **Automatic**

**CorrectIlluminationApply**

This module applies the illumination function created in the previous step to correct the GFP channel images. The panel settings are as follows:

- Select the input image **GFP_ILLUM**
- Name the output image **GFP_ILLUM_CORRECTED**
- Select the illumination function **GFP_ILLUM**
- Select how the illumination function is applied **Subtract**

**IdentifyPrimaryObjects**

This module identifies biological objects of interest and requires grayscale images containing bright objects on a dark background. The nuclei were identified as the primary objects. The panel settings are as follows:

- Use Advanced settings?: **Yes**
- Select the input image: **DAPI_ILLUM_CORRECTED**
- Name the primary objects to be identified: **Nuclei**
- Typical diameter of objects, in pixel units (Min,Max): **5, 40**
- Threshold strategy: **Adaptive**
- Thresholding method: **Otsu**
- Two-class or three-class thresholding: **Three classes**
- Assign pixels in the middle intensity class to the foreground or the background: **Foreground**
- Threshold smoothing scale: **1.3488**
- Threshold correction factor: **0.79**
- Size of adaptive window: **48**
- Automatically calculate size of smoothing filter for declumping: **No**
- Size of smoothing filter: **6**
- Speed up by using lower-resolution image to find local maxima?: **No**

The thresholding strategy, thresholding method, threshold smoothing scale, threshold correction factor, size of the adaptive window, and size of the smoothing filter may vary across images. It is best to maintain consistent conditions during the imaging step to prevent the need to change the settings for every image. The module calls a new widget that displays the primary objects identified. The image histogram was examined (right-click and “Show image histogram”) to approximate the threshold. The value at the elbow of the curve was used and the threshold correction factor was calculated as (New Threshold/ Current Threshold). The segmentation was examined and if the threshold was too sensitive or not sensitive enough, the process was repeated with a new threshold value.

**EditObjectsManually**

This module creates, removes and edits objects previously defined. If the noise around the corners of the images worsened the segmentation in those areas, the nuclei in those areas were deleted. The panel settings are as follows:

- Select the objects to be edited: **Nuclei**
- Name the edited objects: **Nuclei_First_edit**
- Numbering of the edited objects: **Retain**
- Display a guiding image?: **Yes**
- Select the guiding image: **GFP_ILLUM_CORRECTED**

The module calls a new widget where the objects are edited. The nuclei were examined for proper and complete segmentation. The guiding image was used to merge closely associated nuclei if they were determined to be part of one cell.

**IdentifySecondaryObjects**

This module identifies objects using other objects as a starting point. The cytoplasms were identified as secondary objects using their corresponding nuclei. The panel settings are as follows:

- Select the input image: **GFP_ILLUM_CORRECTED**
- Select the input objects: **Nuclei_First_Edit**
- Name the objects to be identified: **Cytoplasm**
- Select the method to identify the secondary objects: **Watershed-Image**
- Threshold strategy: **Global**
- Thresholding method: **Otsu**
- Two-class or three-class thresholding: **Three classes**
- Assign pixels in the middle intensity class to the foreground or the background: **Background**
- Threshold smoothing scale: **1.3488**
- Threshold correction factor: **1.2**
- Log transform before thresholding? : **Yes**
- Discard secondary objects touching the border of the image? : **Yes**
- Discard the associated primary objects touching the border of the image? : **Yes**
- Name the new primary objects: **Nuclei_After_Secondary_Object**

The same threshold calculation method was used as for the primary objects.

**MeasureObjectSizeShape**

This module measures several area and shape features of identified objects. Several area and shape features were taken of cytoplasms and nuclei . The panel settings are as follows:

- Select object sets to measure: **Cytoplasm** and **Nuclei_After_Secondary_Object**

**CalculateMath**

This module takes measurements produced by previous modules and performs basic arithmetic operations. The area of the cytoplasm objects was divided by the area of their corresponding nucleus objects. The settings for the panel are as follows:

- Name the output measurement : **Cytoplasmic_Nuclear_Ratio**
- Operation : **Divide**
- Select the numerator measurement type : **Object**
- Select the numerator objects: **Cytoplasm**
- Select the numerator measurement
  - Category: **AreaShape**
  - Measurement: **Area**
- Select the denominator measurement type: **Object**
- Select the denominator objects: **Nuclei_After_Secondary_Object**
- Select the denominator measurement
  - Category: **AreaShape**
  - Measurement: **Area**

**FilterObjects**

This module removes selected objects based on measurements produced by another module. The nuclei were filtered according to the area covered by their corresponding cytoplasms. The nuclear area had to be at least 1.5x smaller than the cytoplasmic area or else it was discarded. The settings for the panel are as follows:

- Select the objects to filter: **Nuclei_After_Secondary_Object**
- Name the output objects: **FILTERED_NUCLEI**
- Select the filtering mode: **Measurements**
- Select the filtering method : **Limits**
- Select the measurement to filter by:
  - Category: **Math**
  - Measurement: **Cytoplasm_nuclei_ratio**
- Filter using a minimum measurement value? : **Yes**
- Minimum value: **1.50**
- Filter using a maximum measurement value? : **No**

The minimum value can be changed to be more or less stringent with the filtering.

**FilterObjects**

The cytoplasms were filtered according to the area they covered in respect to their nuclei. The cytoplasmic area had to be at least 1.5x larger than nuclear area or else it was discarded. The panel settings are as follows:

- Select the objects to filter: **Cytoplasm**
- Name the output objects: **FILTERED_Cytoplasm**
- Select the filtering mode: **Measurements**
- Select the filtering method : **Limits**
- Select the measurement to filter by:
  - Category: **Math**
  - Measurement: **Cytoplasm_nuclei_ratio**
- Filter using a minimum measurement value?: **Yes**
- Minimum value: **1.50**
- Filter using a maximum measurement value?: **No**

**RelateObjects**

This module assigns a relationship to objects so that nuclei can be grouped with their cytoplasms. The panel settings are as follows:

- Parent objects: **FILTERED_Cytoplasm**
- Child Objects: **FILTERED_NUCLEI**
- Do you want to save the children with parents as a new object set? **Yes**
- Name the output object: **RELATED_CYT_NUC**

**MaskObjects**

This module removes objects outside of a specified region. The nuclei mask was used in conjunction with the cytoplasm mask to create a mask that contained the cytoplasms excluding the area covered by the nuclei. The settings for the panel are as follows:

- Select objects to be masked: **FILTERED_Cytoplasm**
- Name the masked objects: **Cytoplasm_Without_Nuclei**
- Mask using a region defined by other objects or by binary image: **Objects**
- Select the masking object: **FILTERED_NUCLEI**
- Invert the mask?: **Yes**
- Handling of objects that are partially masked: **Keep overlapping region**
- Numbering of resulting objects: **Retain**

**MeasureObjectIntensity**

This module measures several intensity features for the identified objects. These measurements were taken using the cytoplasmic (excluding nuclei) and filtered nuclei masks on the GFP channel images. The settings for the panel are as follows:

- Select Images to measure: **GFP** and **GFP_ILLUM_CORRECTED**
- Select objects to measure: **Cytoplasm_Without_Nuclei** and **FILTERED_NUCLEI**

**MeasureObjectIntesityDistribution**

This module measures the spatial distribution of intensities within each object. These measurements were taken using the cytoplasmic (excluding nuclei) and filtered nuclei masks on the GFP channel images. The settings for the panel are as follows:

- Calculate intensity Zernikes?: **Magnitudes and phase**
- Select images to measure: **GFP** and **GFP_ILLUM_CORRECTED**
- Select objects to measure: **Cytoplasm_Without_Nuclei**
- Object to use as center? **Edges of other objects**
- Select objects to use as centers: **FILTERED_NUCLEI**
- Number of bins: **6**

**CalculateMath**

The Integrated Intensity of the cytoplasm was divided by the integrated intensity of the nucleus. The settings for the panel are as follows:

- Name the output measurement : **CDK2_CYT_NUC_RATIO**
- Operation : **Divide**
- Select the numerator measurement type : **Object**
- Select the numerator objects: **Cytoplasm_Without_Nuclei**
- Select the numerator measurement
  - Category: **Intensity**
  - Measurement: **IntegratedIntensity**
  - Image: **GFP_ILLUM_CORRECTED**
- Select the denominator measurement type: **Object**
- Select the denominator objects: **FILTERED_NUCLEI**
- Select the denominator measurement
  - Category: **Intensity**
  - Measurement: **IntegratedIntensity**
  - Image: **GFP_ILLUM_CORRECTED**

**MeasureObjectSizeShape**

Several area and shape features were taken of the filtered cytoplasmic (excluding nuclei) and filtered nuclei masks . The panel settings are as follows:

- Select object sets to measure: **Cytoplasm_Without_Nuclei** and **FILTERED_NUCLEI**

**ExportToSpreadsheet**

This module exports measurements into one or more files that can be opened in Excel or other spreadsheet programs. The panel settings are as follows:

- Filename prefix: **{experiment#}**
- Select the measurements to export: **Yes**
- Press button to select measurements: **Cytoplasm_without_nuclei** and **FILTERED_NUCLEI**

The ouput folder was specified in the “Output Settings” tab. The relevant files used were the “{experiment#}_Cytoplasm_Without_Nuclei.csv**”** and “{experiment#}_FILTERED_NUCLEI.csv” files. The files were imported into R studio and the Intensity_IntegratedIntensity_GFP_ILLUM_CORRECTED values from each file and the Math_CDK2_CYT_NUC_RATIO was extracted into one data frame. The data frame was filtered to remove all rows where the cytoplasmic integrated intensity and the nuclear integrated intensity were both below 1.1, the nuclear integrated intensity was less than 1.1 while the cytoplasmic integrated intensity was greater than 1.1, and where the cytoplasmic integrated intensity was below 1.1. This was done to remove rows where negligible integrated intensity values were producing outliers in the cytoplasmic/nuclear ratio.

CellProfiler requires the following citations to be used in the paper but I think they said its okay to only use the most recent one:

- Stirling DR, Swain-Bowden MJ, Lucas AM, Carpenter AE, Cimini BA, Goodman A (2021). CellProfiler 4: improvements in speed, utility and usability. BMC Bioinformatics, 22 (1), 433. . PMID: 34507520 PMCID: PMC8431850. [doi](https://doi.org/10.1186/s12859-021-04344-9). [pdf](https://cellprofiler.org/files/cellprofiler/files/153_Stirling_BMCBioinf_2021.pdf).
- McQuin C, Goodman A, Chernyshev V, Kamentsky L, Cimini BA, Karhohs KW, Doan M, Ding L, Rafelski SM, Thirstrup D, Wiegraebe W, Singh S, Becker T, Caicedo JC, Carpenter AE (2018). CellProfiler 3.0: Next-generation image processing for biology. PLoS Biol. 16(7):e2005970. PMID: 29969450. [doi](https://doi.org/10.1371/journal.pbio.2005970).
- Kamentsky L, Jones TR, Fraser A, Bray M, Logan D, Madden K, Ljosa V, Rueden C, Harris GB, Eliceiri K, Carpenter AE (2011). Improved structure, function, and compatibility for CellProfiler: modular high-throughput image analysis software. Bioinformatics 2011. PMID: 21349861 PMCID: PMC3072555. [doi](http://bioinformatics.oxfordjournals.org/content/27/8/1179.full.pdf?keytype=ref&ijkey=ujLFUXwONdtX58c).
- Carpenter AE, Jones TR, Lamprecht MR, Clarke C, Kang IH, Friman O, Guertin DA, Chang JH, Lindquist RA, Moffat J, Golland P, Sabatini DM (2006) CellProfiler: image analysis software for identifying and quantifying cell phenotypes. Genome Biology 7:R100. PMID: 17076895 [[link to paper at Genome Biology](http://genomebiology.com/2006/7/10/R100)
